# Mitochondrial inner membrane remodeling as a driving force of organelle shaping

**DOI:** 10.1101/2024.05.27.595707

**Authors:** Noga Preminger, Ben Zucker, Sarah Hassdenteufel, Till Stephan, Stefan Jakobs, Michael M Kozlov, Maya Schuldiner

## Abstract

Mitochondria are dynamic organelles exhibiting diverse shapes. While the variation of shapes, ranging from spheres to elongated tubules, and the transition between them, are clearly seen in many cell types, the molecular mechanisms governing this morphological variability remain poorly understood. Here, we propose a novel shaping mechanism based on the interplay between the inner and outer mitochondrial membranes. Our biophysical model suggests that the difference in surface area, arising from the pulling of the inner membrane into cristae, correlates with mitochondrial elongation. Analysis of live cell super-resolution microscopy data supports this correlation, linking elongated shapes to the extent of cristae in the inner membrane. Knocking down cristae shaping proteins further confirms the impact on mitochondrial shape, demonstrating that defects in cristae formation correlate with mitochondrial sphericity. Our results suggest that the dynamics of the inner mitochondrial membrane are important not only for simply creating surface area required for respiratory capacity, but go beyond that to affect the whole organelle morphology. This work explores the biophysical foundations of individual mitochondrial shape, suggesting potential links between mitochondrial structure and function. This should be of profound significance, particularly in the context of disrupted cristae shaping proteins and their implications in mitochondrial diseases.

## Introduction

Mitochondria, highly dynamic and shape-shifting organelles, display remarkable morphological variations in diverse organisms, cell types, metabolic states, and even within a single cell (Kuznetsov *et al*, 2006a; Kuznetsov & Margreiter, 2009). While the basic unit of mitochondria is conventionally considered as the tubular, “bean-like” structure observed in electron micrographs and portrayed in textbooks, the shape of an individual mitochondrion is more intricate than conventionally depicted. A single mitochondrion can manifest a wide spectrum of shapes, spanning from a spherical shape through elongated tubular structures, and even more complex shapes such as donuts and lassos (Preminger & Schuldiner, 2024; Place *et al*, 2023). Spherical mitochondria are evident in pathological states, often seen as mitochondrial swelling, but they also occur in healthy states within specific cell types, like hepatocytes (Das *et al*, 2012a), or within morphologically heterogeneous mitochondrial populations within a single cell (Kuznetsov *et al*, 2006b). In contrast, on the tubular side of mitochondrial shapes, mitochondria can generate lengthy, narrow tubules extending several microns within the cell (Rafelski, 2013).

Mitochondria are structurally distinct from other organelles. The lumen of a mitochondrion is enclosed by a double membrane structure consisting of the outer mitochondrial membrane (OMM) and the inner mitochondrial membrane (IMM). The IMM itself can be divided to two distinct subdomains: the inner boundary membrane (IBM), which runs parallel to the OMM, and intricate membrane invaginations known as cristae, which function as the main sites of energy conversion in mitochondria (Cogliati *et al*, 2016). Cristae, resembling thin membrane cisternae, extend into the mitochondrial lumen and are connected to the IBM by narrow membrane bridges known as cristae junctions (CJs). The IMM is tightly connected to the OMM in contact sites mediated by the mitochondrial intermembrane space bridging (MIB) complex (Ott *et al*, 2012; Tang *et al*, 2019). The MIB complex is formed by the sorting and assembly machinery (SAM) complex in the outer membrane and the mitochondrial contact site and cristae organizing system (MICOS) in the inner membrane. MICOS plays a major role in CJ formation and maintenance (Stephan *et al*, 2020). The CJs and the cristae are dynamic structures that undergo constant rearrangements and remodeling in response to metabolic and physiological adaptations (Kondadi *et al*, 2020; Stephan *et al*, 2019; Cogliati *et al*, 2016).

Cristae architecture is governed by the interplay of MICOS and several other players, among them the IMM protein Optic atrophy 1 (OPA1). OPA1 has a role in mitochondrial fusion and fission as well as cristae organization (Cipolat *et al*, 2004; MacVicar & Langer, 2016). The latter is due to OPA1 maintaining a negative membrane curvature and controlling the width of the CJ (Anand *et al*, 2021).

Depletion of OPA1 leads to a dramatic change in cristae morphology, although it is not essential for the formation of CJs (Barrera *et al*, 2016). Additionally, the F_1_F_0_-ATP synthase, besides its well-known enzymatic activity as the last step of the respiration pathway, is also a pivotal contributor to cristae structure. The F_1_F_0_-ATP synthase organizes into ribbon-like rows of dimers along cristae ridges and induces high membrane curvature at the rim and tip of cristae (Strauss *et al*, 2008; Davies *et al*, 2012; Blum *et al*, 2019).

While extensive data have been accumulated on the proteins involved in cristae formation and shaping, the mechanisms responsible for shaping of the OMM, and hence a mitochondrion as a whole, remain largely elusive. Unlike many other tubular structures in the cell, such as the endoplasmic reticulum, T-tubules and intracellular transport intermediates (Kawaguchi & Fujita, 2024; Wang *et al*, 2021; Polishchuk *et al*, 2009), little is currently known about protein-based machinery that could control the shaping of mitochondria and the shift between spherical and tubular shapes independently of fission and fusion (Preminger & Schuldiner, 2024). Recent years have brought about a barrage of mechanistic studies on proteins important for fission and fusion, and even though these processes explain a considerable portion of mitochondrial shapes and morphological changes, they fall short of fully explaining the diverse array of shapes observed in individual mitochondria. Despite the prevalence of the morphological diversity at the level of the individual mitochondrion, the molecular and biophysical mechanisms underlying the formation of tubules from spheres and the regulation of these transitions independently of fission and fusion, remain unknown. This limits our capacity to experimentally control mitochondrial shape by loss/gain of function experiments and test the functional significance of their structural diversity.

Here, we propose a novel mechanism for the transition between spherical and elongated shapes of mitochondria, based on the mechanical interplay between the two mitochondrial membranes. We suggest that the driving force for elongation of an individual mitochondrion originates from the area difference between the OMM and the IBM, a difference that arises from the transition of a portion of the IBM area into newly formed cristae, or *de novo* cristae formation. Therefore, our model suggests a correlation between the extent of the mitochondrion shape elongation and the overall area of the cristae. Analysis of live cell super-resolution data demonstrates a clear relationship between the degree of sphericity of mitochondria and the amount of IMM incorporated into cristae. We further support the model by demonstrating that knockdowns of cristae shaping proteins and disruption of cristae formation lead to observable changes in mitochondrial shapes. These findings propose a new way by which regulation of *de novo* cristae formation can govern mitochondrial shape with possible implications for aging, neurodegeneration and mitochondrial diseases.

## Results

### Modelling predicts cristae formation as a driving force for organelle elongation

To support the proposed mechanism of mitochondrial elongation based on the interplay between the mitochondrial membranes, we developed and analyzed a theoretical model. This model considers a system composed of two closed membranes – the outer membrane describes the OMM and the inner represents the IBM. The membranes are parallel to each other with a fixed distance, d, between them (Figure 1A). This distance is maintained by interactions between the two membranes, such as the MICOS-supported contacts between the membranes. Notably, this model is simplistic in that it does not account for a possible addition or removal of lipids to or from the system, which does occur in biological systems, albeit maybe less so in small mitochondria with limited capacity to form contact sites.

**Figure 1.**
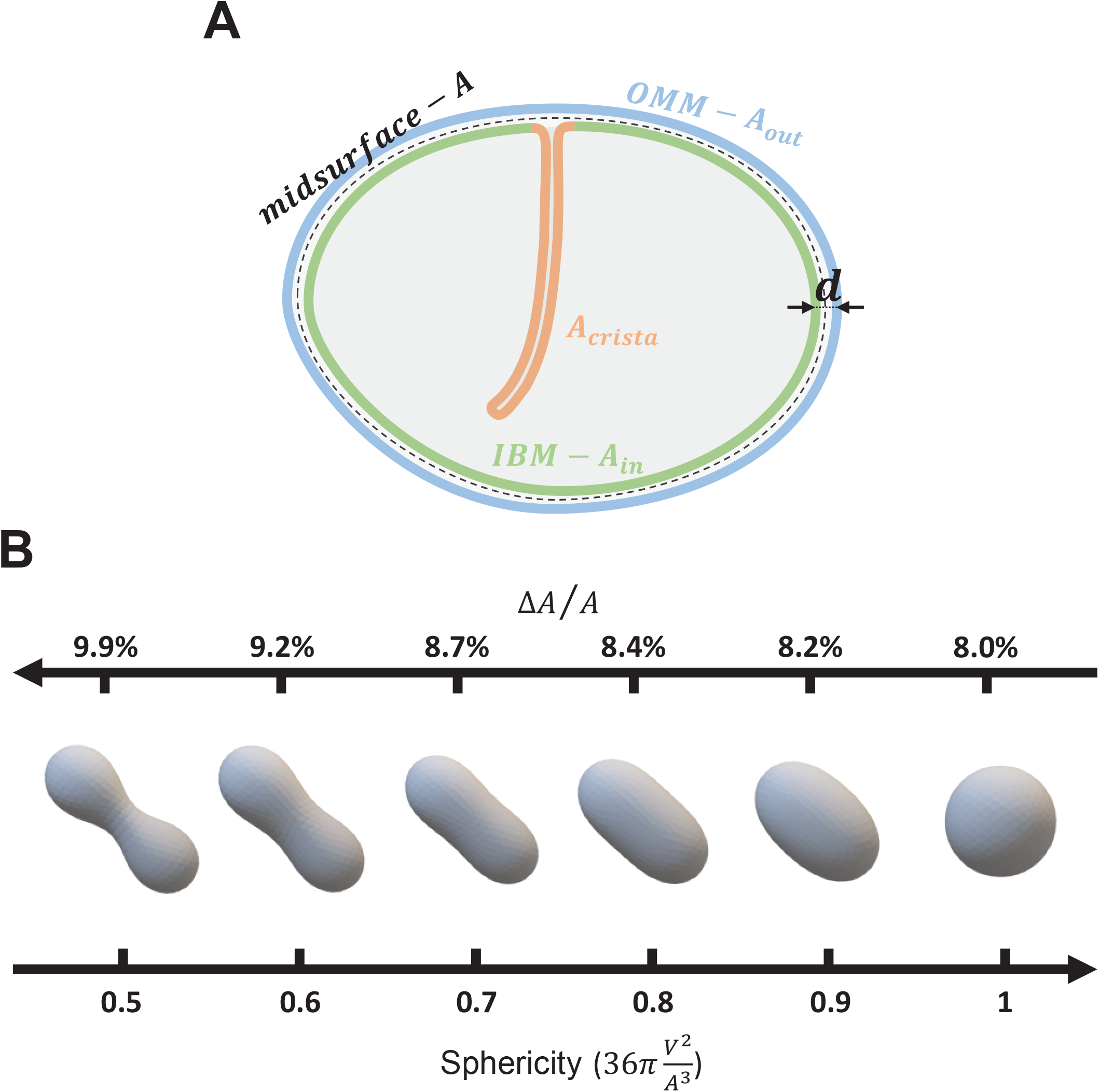
Modelling membrane surface area as a driving force for organelle shape change. (A) A schematic representation and main parameters of the model. The OMM is shown by the blue curve and its surface area denoted by *A*_*OUt*_. The IMM is composed of (i) the IBM, which is shown by the green curve and its surface area denoted by *A*_*in*_, and (ii) the cristae membrane, which is shown by the orange curve. The mid-surface, which is the imaginary surface found between the outer and inner membranes, is shown by the dashed curve and its surface area denoted by A. The distance between the inner and outer membranes is d. (B) Computed results of elastic energy minimization of shapes with a varying difference between the inner and outer membrane surface area. Starting from a spherical shape, as the area difference increases, the sphericity decreases. At small perturbations, the sphere elongates and resembles a prolate ellipsoid. With further increased area difference, the middle part of the shape constricts in a peanut-like structure. The shapes normalized area differences, LJA/A, are calculated for shape evolution starting with a sphere with a diameter of 500 nm and intermembrane distance d= 10 nm, while preserving the surface area during the shape evolution.

Our model is based on a well-established geometrical principle that governs the shape of a sandwich-like system comprised of two closed parallel surfaces. According to this principle, the relationship between the areas of these surfaces dictates the overall shape of the system. Specifically, if the area of the inner surface, *A*_*in*_, is smaller than that of the outer surface, *A*_*out*_, the system tends to adopt an overall elongated (prolate) shape, whereas if *A*_*in*_ increases relative to *A*_*out*_, the shape flattens and becomes more spherical (becomes oblate and biconcave (Seifert, 1997)). This principle underlies the Bilayer Couple Model of a single membrane, where the two leaflets of the membrane can adopt different surface areas due to variations in their lipid and/or protein content (Seifert, 1997; Sheetz & Singer, 1974; Svetina & Žekš, 1989). Here, we use this principle to propose that the elongation of a mitochondrion is driven by a decrease in the IBM area, *A*_*in*_, compared to that of the OMM, *A*_*out*_, due to the folding of the former into new cristae. We termed this concept the CORSET model, which stands for Coupled Outer-inner membrane Rearrangement for Shape Elongation Transition.

To support this proposal quantitatively, we consider the following geometrical and physical arguments (see Materials and Methods for full details). We describe the system’s shape by that of an imaginary mid-surface of area, A, which lies between the OMM and IBM parallel to and at a distance d/2 from each of them (Figure 1A). The shape of the mid-surface is determined at each point by the mean curvature, J. The overall mid-surface’s shape is characterized by the value of J averaged over the surface area, ⟨*J*⟩. The larger the average curvature, ⟨*J*⟩, the more elongated and tubular the system’s shape.

The value of the average curvature, ⟨*J*⟩, is determined by: 1) the difference between the outer- and inner-membrane areas, Δ*A = A*_*out*_ *-A*_*in*_; 2) the inter-membrane distance, d; and 3) the mid-surface area, *A*. A larger ratio 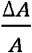 corresponds to a greater average curvature ⟨*J*⟩, and, hence, a stronger elongation of the system’s shape.

To illustrate the effect of the variations of the area difference, Δ*A*, on the system’s shape we performed a computation analogous to those of (Seifert *et al*, 1991), whose essence was a minimization of the bending elastic energy of the system and identifying the optimal shapes that minimize it for different values of 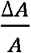 (see Materials and Methods). The extent of elongation can be described by simple geometrical measures, such as the sphericity of the shapes, defined as the ratio between the shape’s squared volume and its cubed surface area, normalized by this ratio’s value corresponding to that of a sphere. Hence, a perfect sphere has a sphericity of one. Sphericity decreases as the shape elongates (Figure 1B). As expected, an increase in Δ*A* drives shape elongation (Figure 1B). Larger 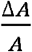 results in significant elongation and leads to substantial deviations from ellipsoidal shapes towards peanut-like shapes with a slightly constricted middle part (Figure 1B, left shape).

### The extent of IMM pulled into cristae correlates with tubularity

The CORSET model predicts that for the initial transition from a spherical mitochondrion to a more tubular shape, more IBM has to be pulled into new cristae. To test this prediction, we assayed whether the shape of mitochondria correlates with the extent of cristae. Since this exact comparison is, in fact, technically challenging, we decided to use as a proxy the percentage of cristae surface area out of the total surface area of the IMM (which includes both the IBM and the cristae, from here on called [cristae/IMM]%). To do that, we analyzed images of mitochondria from live HeLa cells acquired by 2D live-cell stimulated emission depletion (STED) microscopy, in which the entire IMM was stained with the mitochondrial dye PK Mito Orange (PKMO) (Liu *et al*, 2022). Mitochondrial shape was traced and assessed for its sphericity (Figure 2A). To determine sphericity, we calculated the two-dimensional circularity, defined as 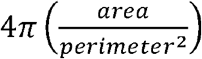 . Indeed, when [cristae/IMM]% was calculated, we observed a correlation between circularity and the percentage of IMM incorporated into cristae, indicating that more spherical mitochondria had lower [cristae/IMM]% (Figure 2B). A similar trend was observed when the calculations were transformed from two-dimensional to an inferred three-dimensional prediction, considering the surface area of disk-shaped cristae and the surface area of the cylinder-shaped boundary membrane (see Supplementary figure 1A and B).

**Figure 2.**
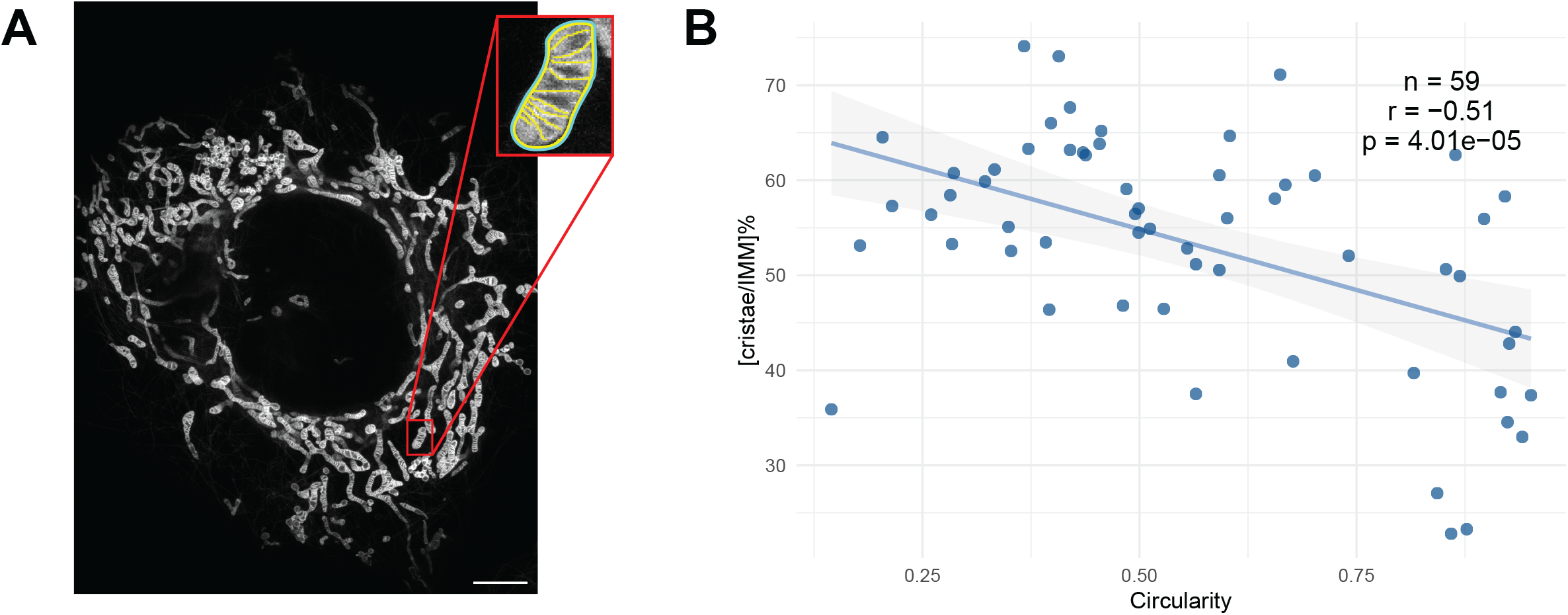
The extent of cristae correlates with degree of circularity. (A) HeLa cells were stained with PKMO (inner membrane dye) and imaged using live-cell STED. The IBM and cristae of individual mitochondria (example in inset) were traced and measured. Scale bar: 5µm (B) The percentage of cristae surface area out of the total surface of the IMM in each mitochondrion ([cristae/IMM]%) was plotted against the circularity measured for the shape of the mitochondrion, where a circularity value of 1 indicates a perfect circle. Data information: n – number of analyzed mitochondria; r – Pearson’s correlation; p – P-value.

Our model is relevant only to the primary shape change – from sphere to an elongated shape, but does not account for further extension of larger, longer tubules. We therefore examined the relationship between shape circularity and [cristae/IMM]% among different size-groups of mitochondria, which could indicate whether they are in the relevant initial shaping stage or have already progressed beyond it. To discern distinct shaping behaviors, the pool of all mitochondria was divided based on the area of the mitochondrion (measured as the two-dimensional area found inside the perimeter of the IBM), using the median value to create two groups (designated as “small” and “large” based on their respective areas) (Supplementary figure 2A). The calculated [cristae/IMM]% was then plotted against the circularity (Supplementary figure 2B). Opposite trends were observed in the small- and large-sized mitochondria groups: while the small-sized mitochondria showed a negative correlation between [cristae/IMM]% and circularity, as expected according to the proposed model, larger mitochondria displayed a positive correlation between [cristae/IMM]% and circularity. These results support a biphasic behavior of the mitochondrial shaping process: an initial shape transition governed by *de novo* cristae formation, followed by further shape extension. During this extension, cristae continue to form to maintain the IBM surface area suitable for respiration, rather than as a means to influence organelle shape. Therefore, we posit that shaping by the interplay of IMM-OMM area difference is relevant to the transition from a perfect sphere to a more elongated shape at earlier stages of the mitochondrion growth, a process that is independent of addition of lipids, but cannot explain the following extensive elongation of mitochondria into long narrow tubules, a stage that requires membrane expansion and increase in lipid mass through fusion or lipid transfer via contact sites.

### Knockdowns of cristae shaping proteins affect mitochondrial shape

If indeed the formation and maintenance of cristae has an impact on the transition from a spherical to elongated shape of the mitochondrion, then changes in the proteins involved in these processes should impact the sphericity of mitochondria in cells. To test this, we knocked down, using siRNA, central proteins involved in cristae formation and maintenance. The MICOS complex, besides its central role in IMM shaping and cristae formation, is also involved in the formation of the mitochondrial intermembrane space bridging (MIB) complex that connects the OMM and IMM (Ott *et al*, 2012; Kozjak-Pavlovic, 2017). Since the integrity of these membrane contact sites is crucial for our model, we did not use MICOS knockdowns that might disrupt these connections, and focused instead on other proteins involved in cristae shaping – OPA1 and the F_1_F_o_□ATP synthase subunit e ATP5ME (Figure 3A, Supplementary figure 3). Electron microscopy (EM) analysis uncovered that, as expected, cristae architecture was altered when knocking down these proteins, resulting in several different cristae morphologies (Figure 3B). We then assessed the shape of mitochondria in these cells, comparing them to mitochondria from control cells. Mean circularity of mitochondria was significantly different between control and knockdown cells (Figure 3C). Knockdowns of OPA1 and ATP5ME resulted in a statistically significant shift in the shape of mitochondria to a more spherical shape as would be predicted by the CORSET model, suggesting that indeed, reduced capacity to form cristae results in less capacity to elongate.

**Figure 3.**
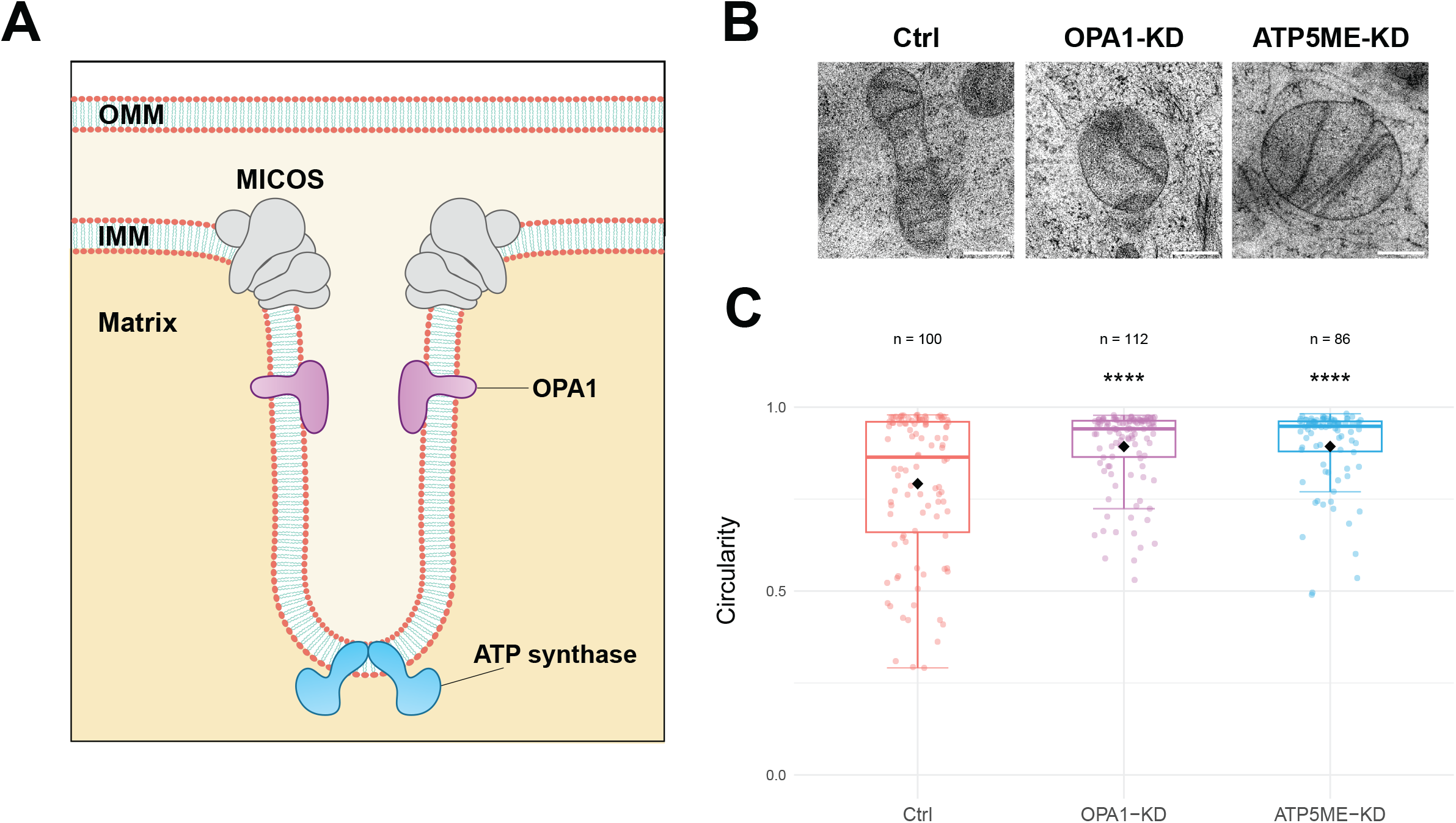
Knockdowns of cristae shaping proteins affect mitochondrial shape. (A) Scheme of a crista and cristae shaping proteins. (B) Representative EM recordings of mitochondria in HeLa cells transfected with a control (Ctrl) siRNA pool or an siRNA pool against cristae shaping proteins, as indicated. Scale bars: 200 nm (C) Circularity measurements of mitochondria sampled from Ctrl and knockdown (KD) cells. Horizontal lines indicate the median, black dots indicate the mean. Student’s t-Test was used to compare to the control. Data information: n – number of analyzed mitochondria; ns (not significant), **p ≤ 0.01, ****p ≤ 0.0001

### Time-resolved shape transitions support a correlation between shape and cristae extent

Examination of still images allowed us to observe a correlation between the overall shape of mitochondria and the proportion of IMM folded into cristae. While static images provide valuable information about the relationship between shape and cristae folding at a specific point, they only offer a snapshot of a dynamic, continuously evolving process. To delve deeper into this interplay, we conducted a time-resolved analysis using previously published time-lapse STED recordings of mitochondria stained with MitoPB Yellow (Wang *et al*, 2019), imaged over prolonged periods. It was shown that in long-term observations (>90s) mitochondria undergo a drastic change in their shape due to photodamage induced by the intense STED lasers. This resulted in a progressive loss of cristae, which gradually retracted into the IBM, and within minutes mitochondria transitioned from narrow tubules to complete spheres (Figure 4A). Importantly, these shape transitions occurred without any fission events. Already this initial description aligns with our model – reduced cristae and increased IBM preceding a change in shape to spherical mitochondria. To validate this we assessed the mitochondrial shape (circularity) and cristae extent ([cristae/IMM]%) at several timepoints during this process. Despite some fluctuations (which might be a by-product of the dynamic cristae moving in and out of the focal plane, thereby somewhat affecting the measurements), an overall reduction in [cristae/IMM]% was observed from the initial tubular shape to the final spherical shape in the majority of mitochondria analyzed (Figure 4B). While the photodamage could induce various changes in mitochondria and consequently in their shape, the association between higher sphericity and a reduction in cristae extent supports our proposed model and highlights the relevance of cristae formation in the shaping of the whole organelle. Analysis of the time-resolved shift in live mitochondria further suggests that the folding or unfolding of cristae may serve as a mechanism to alter surface area of the IBM, potentially contributing to the complex process of mitochondrial shape transition.

**Figure 4.**
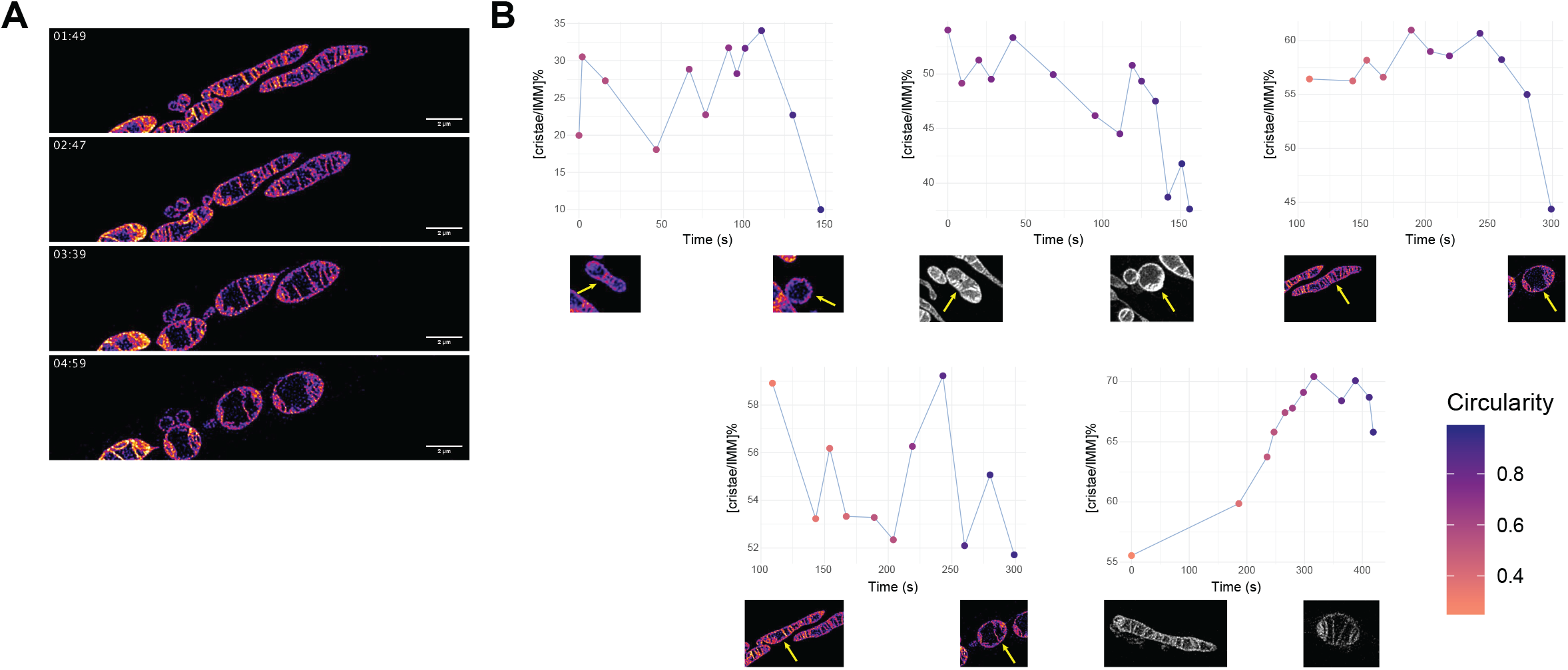
Time-resolved analysis of cristae extent during mitochondrial shape transition. (A) Representative snapshots extracted from time-lapse STED recordings (taken from Wang et al, 2019), showcasing the morphological transformation of mitochondria from tubular to spherical shapes over time. Mitochondria were stained using MitoPB Yellow, enabling visualization of the IMM (Wang et al, 2019). (B) Graphs depicting the change in [cristae/IMM]% over time (seconds). Each graph represents one individual mitochondrion (shown in picture below each graph for initial and final times tracked, arrows indicating the mitochondrion analyzed), with color coding indicating the degree of circularity of the mitochondrial shape at each time point.

## Discussion

Research of mitochondrial shape so far has been dominated by exploration of the network formation with a clear focus on the processes of fission and fusion that are fundamental in shaping it. Fission and fusion, clearly, also affect the length of an individual mitochondrion. However, they cannot fully explain every transition from a sphere to elongated shapes of mitochondria.

In this study, we explored the possible biophysical mechanisms that govern this initial basic stage of mitochondrion shaping. The process of *de novo* cristae formation clearly affects the ultrastructure of the IMM and therefore the surface area supporting the electron transfer chain machinery and consequently cellular respiration, but we suggest that it is also involved in regulating the entire organelle shape, thereby introducing an additional layer of complexity to cristae functionality. Folding of the IMM into cristae results in reduction of the surface area of the IBM, and consequently, increases the difference between the surface area of the IBM and OMM. Based on geometrical considerations and computations of membrane shapes, we propose that the resulting change in the surface area difference, coupled to the strong physical connectivity between the OMM and IBM, can drive the transition of a small isolated mitochondrion from a sphere to an elongated form (Figure 5A). This implies an interplay between the two membranes, where the OMM follows the action of the IMM, contrary to known processes like fission and fusion, where changes typically initiate with the OMM and progress to the IMM. Although current methodologies make it challenging to definitively prove this model, analyses of mitochondrial shapes in live cells support the possibility of this hypothesis, as spherical mitochondria tended to have less of their IMM packed into cristae, relative to more tubular mitochondria in which a larger portion of their IMM was observed in cristae. This was evident in both static images and time-resolved recordings during active shape-changes.

**Figure 5.**
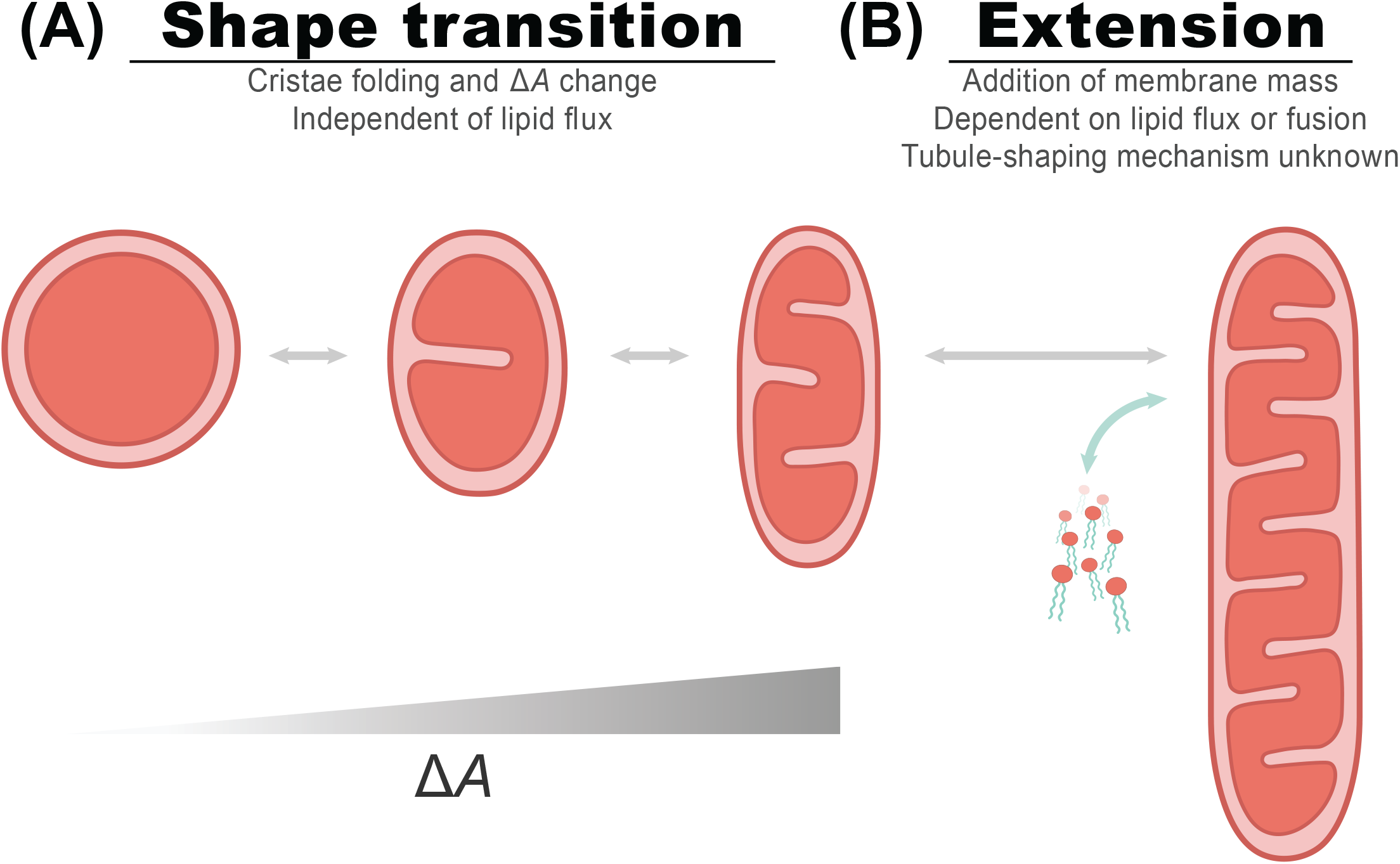
The CORSET model for shaping mitochondria. The shift of mitochondria from spheres to tubules or vice versa can be divided into two substages. (A) The first step is the change from a sphere to a primary tubule (the classical bean-like shape), hypothesized to be driven by cristae formation and area difference changes between the membranes. (B) The second step is a further extension to a full-length tubule by either growth or fusion of several mitochondria, meaning an external addition of lipids, to create longer narrow tubules.

Attempts to validate this theoretical model face various biological and technical constraints. First, the model describes a dynamic membrane remodeling process that occurs in the sub-optical resolution realm and is therefore challenging to track over time with sufficient resolution. In addition, the reliance on 2D measurements to infer 3D shapes introduces inherent limitations in shape descriptors. Consequently, we opted for two complementary methods: super-resolution microscopy and EM. While EM gives outstanding resolution, it relies on thinner slices, making cristae more susceptible to being missed with even slight slice position shifts due to their small size in comparison to the overall organelle shape, which potentially leads to inaccuracies in assessing the cristae extent. Thus, to minimize the possibility of such artifacts, we analyzed cristae extent in super-resolution images of live cells (Jakobs et al, 2020).

Our model describes a shape change that does not require, and even precludes, the addition of lipids to either of the membranes. This assumption is crucial to explain the folding of cristae as a driver of area difference between the membranes. Of course, this also makes it extremely challenging to test in a cell where contact sites are constantly contributing to new lipid uptake by mitochondria (Sassano *et al*, 2022; Scorrano *et al*, 2019). Indeed, in live dynamic mitochondria, lipid addition and loss are constantly occurring, and thus it is important to highlight that our suggested mechanism is suitable for specific cases, for example where the mitochondrial shape has to change without affecting the lipid content or in the absence of functional lipid uptake, and is clearly only one of many processes and mechanisms that help control the shape of mitochondria.

The CORSET model underscores the possible role of *de novo* cristae formation in generation of area difference between the membranes, but it may also involve additional parameters that contribute to shape regulation. For instance, the consistent inter-membrane distance maintained by contact sites is an essential factor for the feasibility of this model. Disruption of this factor could potentially impede the shaping process, serving as another avenue of regulation to this mechanism beyond the cristae shaping machinery. Thus, it is of future interest how changes or regulation of all parameters considered in the model, affect whole mitochondrial morphology.

Analysis of mitochondrial shapes in live cells revealed that the suggested mechanism can only explain a certain population of mitochondria that is defined by their small size. This, together with the underlying assumption of the model for no lipid addition, emphasizes two key points: firstly, the applicability of this model is primarily to small, isolated mitochondria, rather than to elongated tubules or those within the mitochondrial network. Secondly, it highlights the biphasic nature of shape change in mitochondria. The overall shift between a sphere to a long narrow tubule can be divided into two – the first step is the transition from a sphere to a primary tubule (the classical bean-like shape) (Figure 5A), which could be driven completely or mostly by *de novo* cristae formation and change in area difference between the membranes, as explained by our model. Importantly, it is not the mere presence of cristae but rather the folding of new cristae that can drive this process due to the decrease in the surface area of the IBM. The second step of shape shift is the subsequent extension to a full-length tubule by growth or fusion of several mitochondria to create longer tubules (Figure 5B). Elongation at this second step cannot be explained by cristae formation alone, as it is constrained by the lower limit of width of an individual mitochondrion. This lower limit on width represents a physical constraint that impedes additional elongation solely through cristae formation. Essentially, there is a point at which the mitochondrion cannot narrow any further without compromising its structural stability or functional efficiency. In addition, at a certain point expansion of the membranes will require, and even depend on, supplementation of lipids or fusion of entire mitochondrial units. Beyond representation of a substage towards longer tubules, the first step in our model may also allow for a fast and/or transient, short-term shift in shape, without the profound changes and global involvement associated with lipid uptake. Hence, it is essential to distinguish between the substages of shape transition and elucidate the mechanisms, forces, kinetics, and components required for each one.

Spherical mitochondria are observed in various conditions and cell types, both in healthy and pathological states. For example, the spherical mitochondria seen in hepatocytes (Das *et al*, 2012b) or the shapes of mitochondria in skeletal muscles, ranging from small round mitochondria close to the sarcolemma to long tubular mitochondria located at the core of myofibers (Picard *et al*, 2013). While the majority of focus in the literature has been on the pathological state, often referred to as “swelling”, our results suggest that sphericity should be regarded as a normal part of the mitochondria shape repertoire. It is yet unclear whether spherical shapes are just a step on the way to further elongation to a tubule or whether they hold some functional significance. The biphasic model might suggest that mitochondrial fusion is only possible after cristae induced mitochondrial elongation. To study questions regarding the functional importance of spherical and tubular shapes, we must be able to control this shape transition experimentally. However, the regulation that governs the decision on mitochondrial shape and the transition between one shape to another is an unexplored area that requires further research.

The link between mitochondrial diseases and malfunction of cristae shaping proteins has been clearly demonstrated, showing detrimental effects on organellar, cellular and organismal level (Cogliati *et al*, 2016; Colina-Tenorio *et al*, 2020). The impact of disrupted cristae shaping and remodeling has been mainly attributed to dysfunction of respiratory processes that occur in the cristae and are greatly impacted by their shape. We suggest taking into consideration the effect of impaired cristae formation on the overall shape of mitochondria, and to explore the direct implications this could have on mitochondrial and cellular function beyond the clear impact on energy conversion.

## Methods

### Theoretical modelling and analysis

#### Geometrical formulation

To describe the geometry of the system, we considered three mutually parallel surfaces. The outer and inner membranes are represented by their mid surfaces lying between the membrane leaflets and having the areas *A*_*out*_ and *A*_*in*_, respectively (Figure 1A). The system as a whole is described by a surface referred below to as the system surface that is parallel to and lies at half a distance between the surfaces of the outer and inner membranes (Figure 1A). The area of the mid surface is denoted by *A*.

The local shape of the system surface at each point is described by two principal curvatures, *c*_*p1*_ and *c*_*p2*_, which are the curvatures of two arc-like cross-sections of the surface by normal planes in the specific directions called the principal directions (Spivak). The sign of each principal curvature is conventionally defined according to the convexness (positive sign) or concaveness (negative sign) of the corresponding cross-section line. Alternatively, the local surface shape can be described by the sum and product of the principal curvatures, *J*= *c*_*p*1_ + *c*_*p*2_, and *K*= *c*_*p*1_ *· c*_*p*2_ (Helfrich, 1973), referred to as the total and Gaussian curvature, respectively. In the following we will assume that the absolute values of the curvatures, |*c*_*p*1_|, |*c*_*p*2_|, and hence, |*J*| are small all along the system surface compared to the inverse distance between the outer and inner membranes, 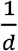, such that their dimensionless combinations are smaller than one, |*c*_*p*1_| *d*« 1, |*c*_*p*2_| *d*« 1, |*J*|d« 1. In this case the dimensionless value of the Gaussian curvature is smaller than that of the total curvature |*K*|*d*^2^ « |*J*|*d*, so that in all expressions below we will neglect the contributions related to K. For brevity, we will refer to *J* as simply the curvature.

Since all three surfaces are mutually parallel, their local geometrical characteristics are interrelated. The area elements, *dA*_*out*_ and *dA*_*in*_, of, respectively, the outer and inner membrane surfaces are related to the element of the system surface, *dA*, by the relationships (see e.g. (Murphy, 1966)),

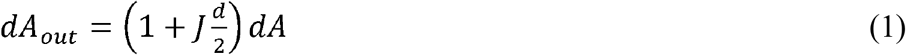

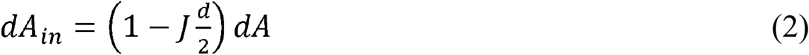

The expressions (Eqs.1,2) enable determination of a qualitative relationship between the overall character of the system shape and the difference between the outer and inner membrane areas, *A*_*out*_, and *A*_*in*_.

Indeed, the outer and inner membrane areas are given by the integration of the area elements, *A*_*out*_ = *∮ dA*_*out*_ and *A*_*in*_ = *∮ dA*_*in*_, so that, according to (Eqs.1,2), they can be expressed by

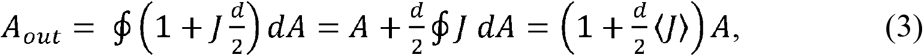

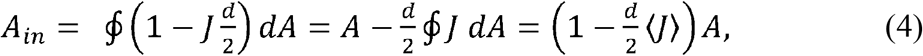

where ⟨*J*⟩ is the curvature of the system surface averaged over the surface area, *A*. Combining (Eqs.3,4) we obtain

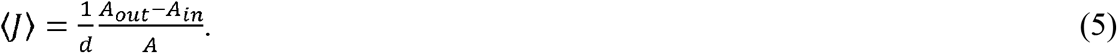

According to (Eq.5), the larger the difference between the outer and inner membrane areas, *A*_*out*_ -*A*_*in*_, the larger is the average curvature of the system surface ⟨*J*⟩.

#### Elasticity analysis

The areas of the outer, *A*_*out*_, and inner, *A*_*in*_, membranes, determine the average curvature, <*J* >, but do not set the exact shape of the system’s surface.

To find the specific shape of the system’s surface, we consider the elastic energy of the system, *F*. The optimal configuration of the system is determined by minimization of the sum of the elastic energies of the outer, *F*_*out*_, an inner, *F*_*in*_, membranes, *F= F*_*out*_ *+ F*_*in*_, by varying the system shape within the constraints imposed by the given areas *A*_*out*_ and *A*_*in*_ of and the given distance, d, between the outer and inner membrane surfaces.

We assume the only contribution to the elastic energy of the membranes to be the bending energy, which implies that the membranes are considered to be non-stretchable. To compute the elastic energies of the outer and inner membranes we use the Helfrich model (Helfrich, 1973) according to which the bending energy of a membrane element related to the membrane unit area depends on the membrane total curvature and can be presented by

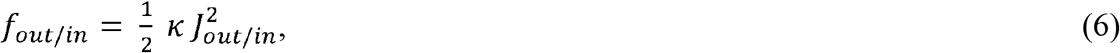

where *κ* is the bending modulus (Helfrich, 1973) and the subscript indicates which membrane is described. The curvature of each membrane could be inferred from the middle surface by geometrical considerations (Murphy, 1966),

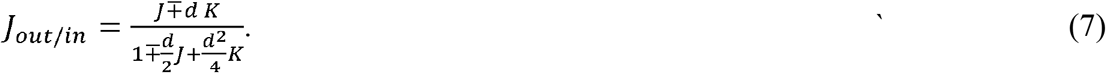

Since, according to (Eq.7), the deviations of *J*_*OUt/in*_ from J are quadratic in the product of the principle curvatures and the membrane thickness, they will be neglected.

The total elastic energy, *F*_*in/OUt*_, is obtained by integrating *f*_*in/OUt*_ over the area of the corresponding membrane surface,

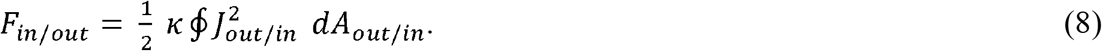

By using the (Eq.6) we assume the membranes to have a vanishing spontaneous curvature (Helfrich, 1973) which implies that for each membrane the two leaflets are similar so that the membrane structure is up-down symmetric.

Such computation of the optimal configuration of our system is technically analogous to the previous analysis of the shapes of individual membranes determined by the Bilayer-Couple model (Svetina & Žekš, 1989; Seifert, 1997).

We numerically performed the minimization of the system bending energy (Eq.8) upon a constraint of constant 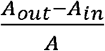 and found the optimal shapes of the system surface by using the specialized program, the Brakke’s Surface Evolver (Brakke, 1992). The results for a few values of the relative difference between the outer and inner membrane areas, 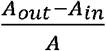, are presented in (Figure 1B). As predicted by the geometrical analysis, an increase in 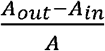 drives the shape elongation. Indeed, the obtained optimal shapes resemble ellipsoids when the deviation from a sphere is small. However, for significant elongation, the shapes deviate substantially from ellipsoids, acquiring a peanut-like shape with a slightly constricted middle part.

### Cell culture for live-cell imaging

HeLa cells were grown in Dulbecco’s Modified Eagle Medium (DMEM) with glutaMAX™ additive and 4.5 g/l glucose (Thermo Fisher Scientific, USA). The culture medium was supplemented with 1% (v/v) penicillin/streptomycin (Sigma Aldrich, Munich, Germany), 1 mM sodium pyruvate (Sigma Aldrich), and 10% (v/v) Fetal Bovine Serum (FBS) (Merck Millipore, Burlington, MA, USA). Cells were cultured in an incubator at 37 °C with 5% CO_2_.

### PKMO labeling for live-cell imaging

HeLa cells were seeded in glass-bottom dishes (ibidi GmbH, Germany) one day prior to imaging. Cells were stained with DMEM containing 350 nM PKMO, 200 nM 4-610CP-CTX (tubulin dye, not shown), and 0.2 μl/ml Quant-iT PicoGreen reagent (mtDNA dye, not shown; Thermo Fisher Scientific) at 37 °C for 40 minutes. Following the staining procedure, cells were washed three times with culture medium and incubated at 37 °C for 60 min to remove unbound dye. The cells were imaged in DMEM buffered by 4-(2-hydroxyethyl)-1-piperazineethanesulfonic acid (HEPES) at room temperature.

### Live-cell imaging

HeLa cells were recorded using a Facility Line microscope (Abberior Instruments) equipped with an Olympus UPlanXAPO 60× oil/NA1.42 objective, a STEDYCON STED microscope (Abberior Instruments) equipped with a CFI Plan Apochromat Lambda D 100× oil/NA1.45 objective, or an Expert Line dual-color STED 775 QUAD scanning microscope (Abberior Instruments) equipped with a UPlanSApo 100×/1.40 Oil [infinity]/0.17/FN26.5 objective. PKMO was excited at 561 nm wavelength, and STED was performed using a pulsed depletion laser at 775 nm wavelength with gating of 1 to 7 ns and dwell times of 5 to 10 μs. Pixel sizes of 25 to 28 nm were used for STED nanoscopy and each line was scanned 3 to 10 times (line accumulations). The pinhole was set to 0.7 to 1.0 AU.

### Image analysis for STED microscopy

PKMO labeled cells were visualized and analyzed using Fiji (version 1.54f). For the analysis, linear mitochondria with well-labeled cristae were chosen (mitochondria that did not have any labeled cristae were excluded from the analysis. To this end, our measurements may be an underestimate to the degree of correlation since we do not have the “no cristae” population which may definitely exist in cells). The boundary shape of mitochondria and individual cristae were manually traced and then measured for their shape descriptors. The measured area of each crista was doubled to account for the fact that each crista consists of two membrane layers. The sphericity of mitochondrial shapes was evaluated by the two dimensional circularity: 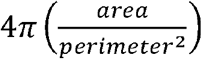.

### Cell culture and transient knockdowns for EM

HeLa cells (ATCC CCL-2) were cultivated at 37 °C in DMEM glutaMAX™, containing 4.5 g/l glucose and 1mM sodium pyruvate (Thermo Fisher Scientific, USA) and supplemented with 10% FBS (Thermo Fisher Scientific, USA) and 1% penicillin/streptomycin (Biological Industries, Israel) in a humidified environment with a 5% CO_2_ atmosphere. For 72h knockdown of MIC60, MIC10, OPA1, or ATP5ME, 2x 10^5^ HeLa cells were seeded into a 60 mm culture dish with regular culture medium. Cells were transfected with the respective high-complexity siRNA pool (siTOOLs Biotech, Germany) to a final concentration of 3 nM using Lipofectamine RNAiMAX transfection reagent (Thermo Fisher Scientific, USA) according to the manufacturer’s instructions. All knockdowns were verified by Western blotting using antibodies against Opa1 (67589, Cell Signaling Technology, USA), Mic10 (ab84969, Abcam, UK), Mic60 (10179-1-AP, Proteintech, USA) and ATPase subunit e (16483-1-AP, Proteintech, USA), respectively.

### EM sample preparation and imaging

EM was performed following a standard sample preparat(Kukulski *et al*, 2012; Bykov *et al*, 2019)et al, 2012; Bykov et al, 2019). In short, a viscous cell suspension was transferred to the 0.1-mm-deep cavity of a 0.1/0.2-mm membrane carrier for a Leica ICE high-pressure freezing machine. The cavity was covered by the flat side of a 0.3-mm carrier, and the sandwich was inserted in the high-pressure freezing machine. Resin embedding was performed using a Leica AFS2 freeze-substitution machine equipped with a processing robot. Samples were embedded in Lowicryl HM20 resin using the freeze-substitution and embedding protocol optimized for in-resin CLEM (Kukulski *et al*, 2012). Dry acetone with 0.1% uranyl acetate was used as the freeze-substitution medium. The blocks were trimmed using a Diatome 45° trimming knife, and 100-nm-thick sections were produced using a Diatome 35° knife on a Leica UC7 microtome. The sections were mounted on 200 mesh copper grids with continuous carbon support film (Electron Microscopy Sciences). Prior to EM, grids were triple post-stained: 1 min in Reynolds lead citrate, 10 min uranyl acetate and 1 min Reynolds lead citrate. EM imaging was performed on a FEI Tecnai G2 F20 TEM microscope operating at 120 kV, equipped with a TVIPS TemCam-XF416 retractable 16-megapixel CMOS camera. SerialEM software was used for data collection (Mastronarde, 2005; Schorb *et al*, 2019), and the detector was operated in the full frame mode (4,096 × 4,096 pixels). To cover a complete cell, a 3 × 3 montage was acquired with a magnification of 6,500× (corresponding to a pixel size of ∼1 nm).

### Image analysis for EM

EM data was visualized and analyzed using Fiji (version 1.54f). Usually, 80-110 mitochondria from at least 8 different cells were randomly selected and analyzed for each sample. The boundary shape of mitochondria manually traced and then measured for its shape descriptors. The sphericity of mitochondrial shapes was evaluated by the two-dimensional circularity: 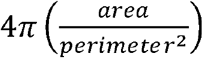 .

## Acknowledgements

We wish to thank Dr. Ofir Klein and Sivan Arad for critical reading of the manuscript and for important feedback. We are grateful to the Weizmann Institute of Science Electron Microscopy unit for their support (and specially to Eyal Shimoni, Nili Dezorella and Sharon Wolf). SJ and MS are supported by an SFB1190 grant from the Deutsche Forschungsgemeinschaft (DFG). SJ is supported by the European Research Council (ERCAdG No. 835102). MS and SH are supported by a grant from The Irving and Cherna Moskowitz Center for Nano and Bio-Imaging (Weizmann Institute of Science). MS and NP are supported by a DFG Middle-Eastern collaboration grant (1028/11-1). MS is an Incumbent of the Dr. Gilbert Omenn and Martha Darling Professorial Chair in Molecular Genetics. MMK is supported by the Israel Science Foundation (grant 1994/22) and holds the Joseph Klafter Chair in Biophysics.

---

**Supplementary figure 1.**
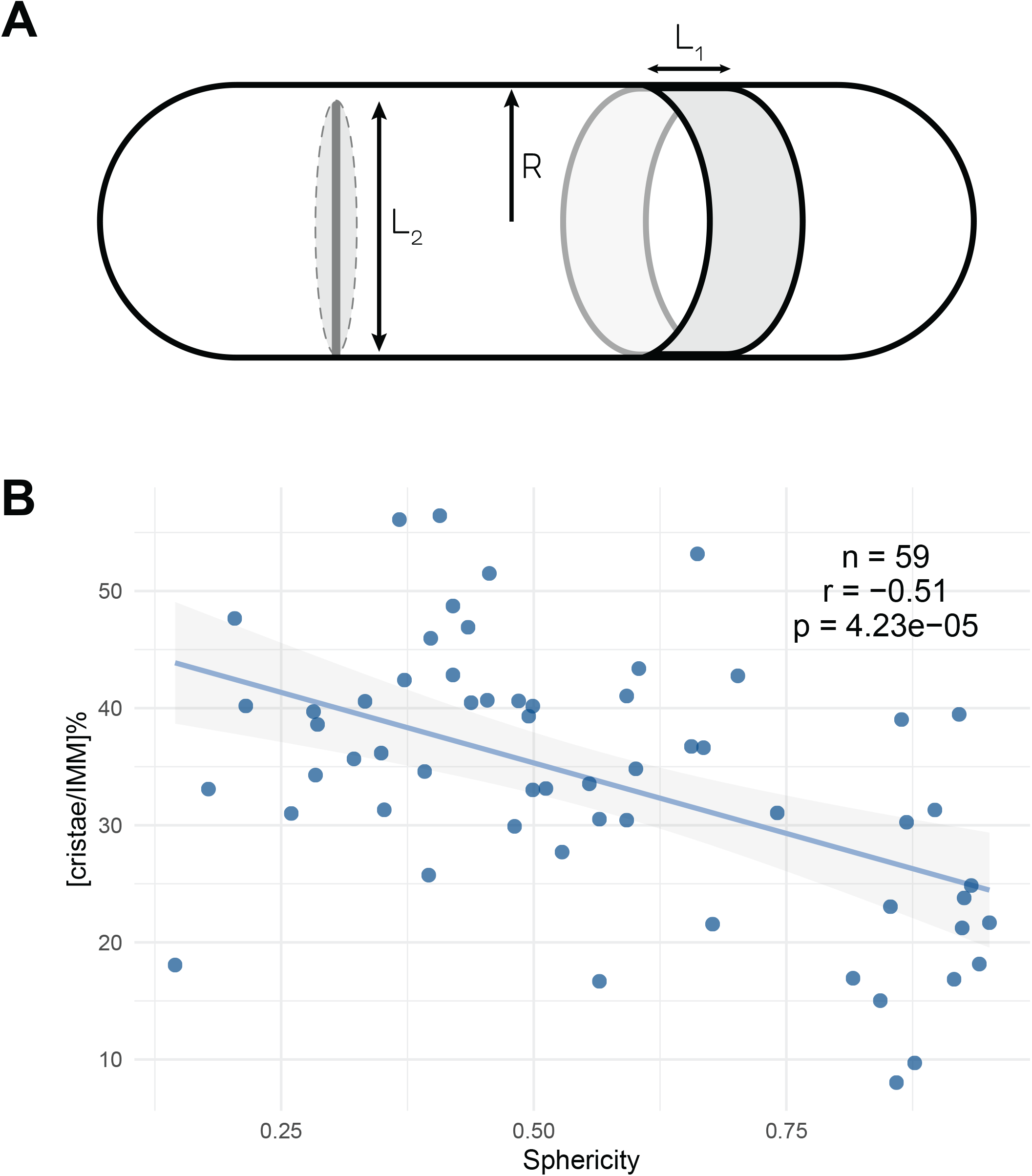
Transformation from two-dimensional (2D) to three-dimensional (3D) calculation maintains a negative correlation between cristae extent and sphericity. (A) A scheme of a mitochondrion in the shape of a cylinder with hemispherical ends, with a radius R. In grey, a ring-shaped portion of the boundary surface area with the width L_1_, and a disk-shaped crista with a diameter of L_2_. For details of 2D-to-3D transformation see Supplementary materials. (B) After estimation of 3D measurements, the percentage of cristae surface area out of the total surface of the IMM in each mitochondrion ([cristae/IMM]%) was plotted against the sphericity measured for the shape of the mitochondrion, where a sphericity value of 1 indicates a perfect sphere. Data information: n – number of analyzed mitochondria; r – Pearson’s correlation; p – P-value.

**Supplementary figure 2.**
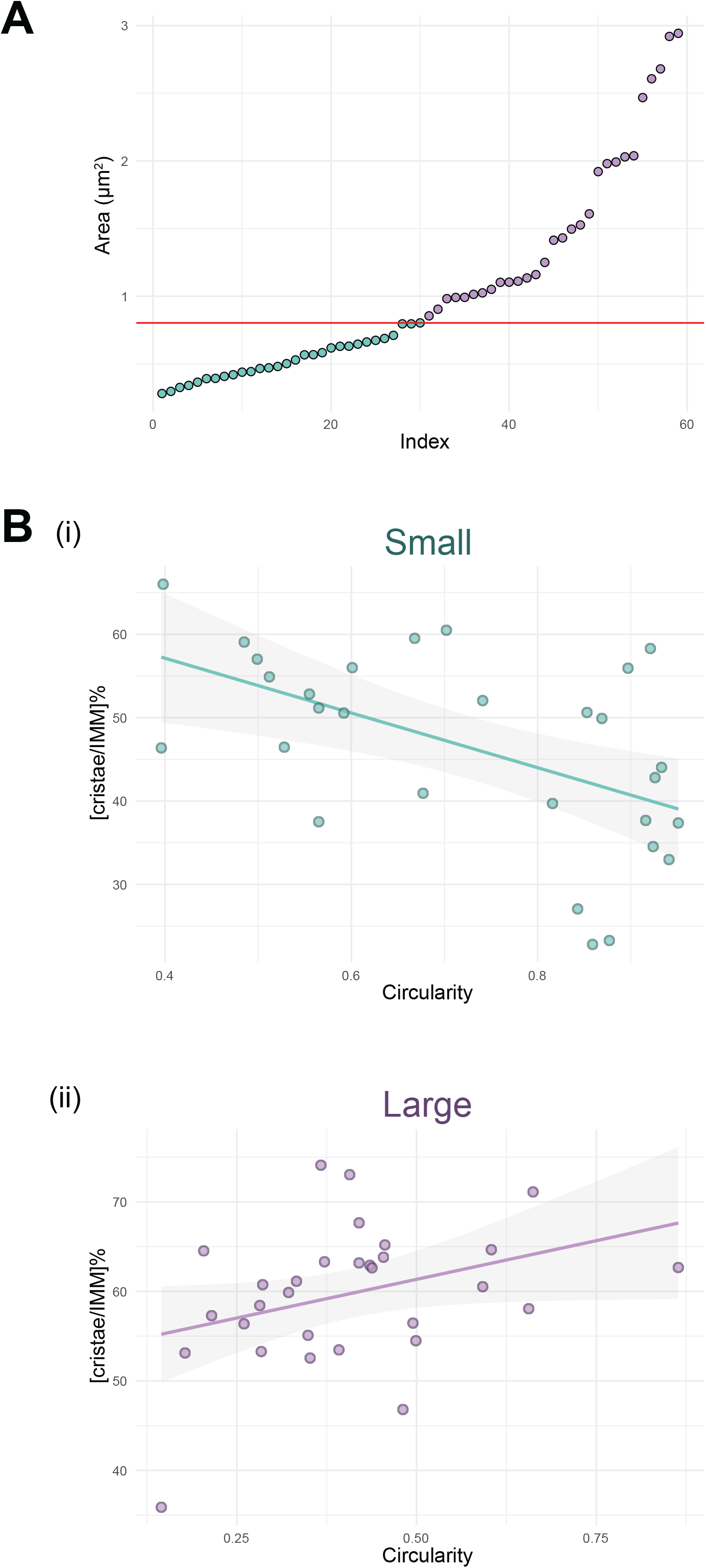
The relationship between [cristae/IMM]% and circularity in different size-groups of mitochondria. (A) Area values (µm^2^) of mitochondria. The data was divided into two size-groups by the median value, indicated by the red line (median = 0.904 µm^2^). (B) The percentage of cristae surface area out of the total surface of the IMM in each mitochondrion ([cristae/IMM]%) was plotted against the circularity measured for the shape of (i) small-sized mitochondria and (ii) large-sized mitochondria. A circularity value of 1 indicates a perfect circle. Data information: n – number of analyzed mitochondria; r – Pearson’s correlation; p – P-value.

**Supplementary figure 3.**
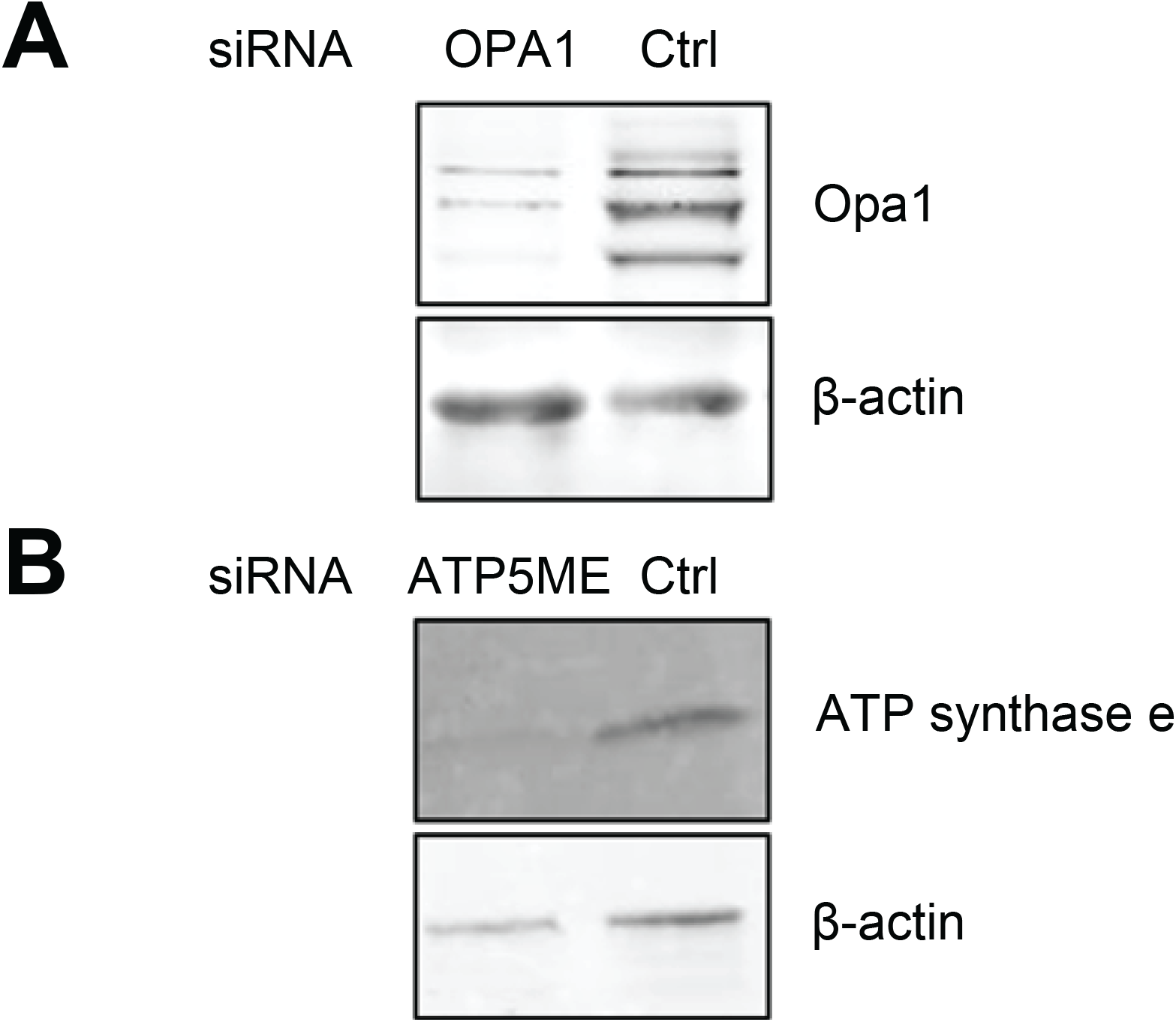
Efficient knockdown of cristae shaping proteins. Western Blots for validation of the siRNA pools used to knockdown OPA1 and ATP5ME. For transfection, 5.2x 10^5^HeLa cells were seeded into a 60mm culture dish with regular culture medium. Upon 72h, cells were harvested and subjected to Western Blotting using the respective antibodies. A) Knockdown of Opa1. B) Knockdown of ATPase subunit e. Beta-actin served as loading control.

## Notes

### Competing Interest Statement

The authors have declared no competing interest.

## References

Anand R, Reichert AS & Kondadi AK (2021) Emerging Roles of the MICOS Complex in Cristae Dynamics and Biogenesis. Biology (Basel) 10

Barrera M, Koob S, Dikov D, Vogel F & Reichert AS (2016) OPA1 functionally interacts with MIC60 but is dispensable for crista junction formation. FEBS Lett 590: 3309–3322

Blum TB, Hahn A, Meier T, Davies KM & Kühlbrandt W (2019) Dimers of mitochondrial ATP synthase induce membrane curvature and self-assemble into rows. Proc Natl Acad Sci U S A 116: 4250–4255

Brakke KA (1992) The Surface Evolver. Exp Math 1: 141–165

Bykov YS, Cohen N, Gabrielli N, Manenschijn H, Welsch S, Chlanda P, Kukulski W, Patil KR, Schuldiner M & Briggs JAG (2019) High-throughput ultrastructure screening using electron microscopy and fluorescent barcoding. Journal of Cell Biology 218: 2797–2811

Cipolat S, De Brito OM, Dal Zilio B & Scorrano L (2004) OPA1 requires mitofusin 1 to promote mitochondrial fusion. Proc Natl Acad Sci U S A 101: 15927–15932

Cogliati S, Enriquez JA & Scorrano L (2016) Mitochondrial Cristae: Where Beauty Meets Functionality. Trends Biochem Sci 41: 261–273

Colina-Tenorio L, Horten P, Pfanner N & Rampelt H (2020) Shaping the mitochondrial inner membrane in health and disease. J Intern Med 287: 645–664

Das S, Hajnóczky N, Antony AN, Csordás G, Gaspers LD, Clemens DL, Hoek JB & Hajnóczky G (2012a) Mitochondrial morphology and dynamics in hepatocytes from normal and ethanol-fed rats. Pflugers Arch 464: 101

Das S, Hajnóczky N, Antony AN, Csordás G, Gaspers LD, Clemens DL, Hoek JB & Hajnóczky G (2012b) Mitochondrial morphology and dynamics in hepatocytes from normal and ethanol-fed rats. Pflugers Arch 464: 101

Davies KM, Anselmi C, Wittig I, Faraldo-Gómez JD & Kühlbrandt W (2012) Structure of the yeast F 1F o-ATP synthase dimer and its role in shaping the mitochondrial cristae. Proc Natl Acad Sci U S A 109: 13602–13607

Helfrich W (1973) Elastic Properties of Lipid Bilayers: Theory and Possible Experiments. Zeitschrift fur Naturforschung - Section C Journal of Biosciences 28: 693–703

Jakobs S, Stephan T, Ilgen P & Brüser C (2020) Light Microscopy of Mitochondria at the Nanoscale. Annu Rev Biophys 49: 289–308

Kawaguchi K & Fujita N (2024) Shaping transverse-tubules: central mechanisms that play a role in the cytosol zoning for muscle contraction. J Biochem 175: 125

Kondadi AK, Anand R, Hänsch S, Urbach J, Zobel T, Wolf DM, Segawa M, Liesa M, Shirihai OS, WeidtkampLJPeters S, et al (2020) Cristae undergo continuous cycles of membrane remodelling in a MICOS LJdependent manner . EMBO Rep 21

Kozjak-Pavlovic V (2017) The MICOS complex of human mitochondria. Cell Tissue Res 367: 83–93

Kukulski W, Schorb M, Welsch S, Picco A, Kaksonen M & Briggs JAG (2012) Precise, Correlated Fluorescence Microscopy and Electron Tomography of Lowicryl Sections Using Fluorescent Fiducial Markers. Methods Cell Biol 111: 235–257

Kuznetsov A V. & Margreiter R (2009) Heterogeneity of Mitochondria and Mitochondrial Function within Cells as Another Level of Mitochondrial Complexity. International Journal of Molecular Sciences 2009, Vol 10, Pages 1911-1929 10: 1911–1929

Kuznetsov A V., Troppmair J, Sucher R, Hermann M, Saks V & Margreiter R (2006a) Mitochondrial subpopulations and heterogeneity revealed by confocal imaging: Possible physiological role? Biochimica et Biophysica Acta (BBA) - Bioenergetics 1757: 686–691

Kuznetsov A V., Troppmair J, Sucher R, Hermann M, Saks V & Margreiter R (2006b) Mitochondrial subpopulations and heterogeneity revealed by confocal imaging: Possible physiological role? Biochimica et Biophysica Acta (BBA) - Bioenergetics 1757: 686–691

Liu T, Stephan T, Chen P, Keller-Findeisen J, Chen J, Riedel D, Yang Z, Jakobs S & Chen Z (2022) Multi-color live-cell STED nanoscopy of mitochondria with a gentle inner membrane stain. Proc Natl Acad Sci U S A 119: e2215799119

MacVicar T & Langer T (2016) OPA1 processing in cell death and disease - the long and short of it. J Cell Sci 129: 2297–2306

Mastronarde DN (2005) Automated electron microscope tomography using robust prediction of specimen movements. J Struct Biol 152: 36–51

Murphy CL (1966) Thermodynamics of low tension and highly curved interfaces. ProQuest Dissertations and Theses

Ott C, Ross K, Straub S, Thiede B, Götz M, Goosmann C, Krischke M, Mueller MJ, Krohne G, Rudel T, et al (2012) Sam50 Functions in Mitochondrial Intermembrane Space Bridging and Biogenesis of Respiratory Complexes. Mol Cell Biol 32: 1173–1188

Picard M, White K & Turnbull DM (2013) Mitochondrial morphology, topology, and membrane interactions in skeletal muscle: a quantitative three-dimensional electron microscopy study. J Appl Physiol 114: 161

Place BC, Troublefield CA, Murphy RD, Sinai AP & Patwardhan AR (2023) Machine learning based classification of mitochondrial morphologies from fluorescence microscopy images of Toxoplasma gondii cysts. PLoS One 18

Polishchuk RS, Capestrano M & Polishchuk E V. (2009) Shaping tubular carriers for intracellular membrane transport. FEBS Lett 583: 3847–3856

Preminger N & Schuldiner M (2024) Beyond fission and fusion – diving into the mysteries of mitochondrial shape. PLoS Biol Accepted

Rafelski SM (2013) Mitochondrial network morphology: Building an integrative, geometrical view. BMC Biol 11: 1–9

Sassano ML, Felipe-Abrio B & Agostinis P (2022) ER-mitochondria contact sites; a multifaceted factory for Ca2+ signaling and lipid transport. Front Cell Dev Biol 10

Schorb M, Haberbosch I, Hagen WJH, Schwab Y & Mastronarde DN (2019) Software tools for automated transmission electron microscopy. Nature Methods 2019 16:6 16: 471–477

Scorrano L, De Matteis MA, Emr S, Giordano F, Hajnóczky G, Kornmann B, Lackner LL, Levine TP, Pellegrini L, Reinisch K, et al (2019) Coming together to define membrane contact sites. Nature Communications 2019 10:1 10: 1–11

Seifert U (1997) Configurations of fluid membranes and vesicles. Adv Phys 46: 13–137

Seifert U, Berndl K & Lipowsky R (1991) Shape transformations of vesicles: Phase diagram for spontaneous-curvature and bilayer-coupling models. Phys Rev A (Coll Park) 44: 1182

Sheetz MP & Singer SJ (1974) Biological Membranes as Bilayer Couples. A Molecular Mechanism of Drug-Erythrocyte Interactions. Proceedings of the National Academy of Sciences 71: 4457–4461

Spivak M A comprehensive introduction to differential geometry vol 2. [PREPRINT]

Stephan T, Brüser C, Deckers M, Steyer AM, Balzarotti F, Barbot M, Behr TS, Heim G, Hübner W, Ilgen P, et al (2020) MICOS assembly controls mitochondrial inner membrane remodeling and crista junction redistribution to mediate cristae formation. EMBO J 39

Stephan T, Roesch A, Riedel D & Jakobs S (2019) Live-cell STED nanoscopy of mitochondrial cristae. Scientific Reports 2019 9:1 9: 1–6

Strauss M, Hofhaus G, Schröder RR & Kühlbrandt W (2008) Dimer ribbons of ATP synthase shape the inner mitochondrial membrane. EMBO Journal 27: 1154–1160

Svetina S & Žekš B (1989) Membrane bending energy and shape determination of phospholipid vesicles and red blood cells. European Biophysics Journal 17

Tang J, Zhang K, Dong J, Yan C, Hu C, Ji H, Chen L, Chen S, Zhao H & Song Z (2019) Sam50–Mic19– Mic60 axis determines mitochondrial cristae architecture by mediating mitochondrial outer and inner membrane contact. Cell Death & Differentiation 2019 27:1 27: 146–160

Wang C, Taki M, Sato Y, Tamura Y, Yaginuma H, Okada Y & Yamaguchi S (2019) A photostable fluorescent marker for the superresolution live imaging of the dynamic structure of the mitochondrial cristae. Proc Natl Acad Sci U S A 116: 15817–15822

Wang N, Clark LD, Gao Y, Kozlov MM, Shemesh T & Rapoport TA (2021) Mechanism of membrane-curvature generation by ER-tubule shaping proteins. Nature Communications 2021 12:1 12: 1–15

