## Supplementary Materials for "Mitochondrial inner membrane remodeling as a driving force of organelle shaping"

### Three-dimensional transformation of shape measurements

To assess the three-dimensional (3D) measurements of analyzed mitochondria from STED images, we estimated the shape of a mitochondrion as a cylinder with hemispherical ends, with a radius of $R$ (Supplementary figure 1A). To assess the 3D surface area of the boundary membrane ($S_{boundary}$) of a mitochondrion, we first considered a ring-shaped portion of the surface area, with a radius of $R$ and a width of $L_{1}$ (Supplementary figure 1A). The surface area of such shape will be

$S=2\pi*R*L_{1}$.

Extending this to the whole boundary membrane (which consists of many such ring-shaped portions), the surface area for each mitochondrion was approximated as

$S_{boundary}=\pi*R*P$,

where $P$ is the perimeter measured by 2D tracing of mitochondria, and $R$ is half of the diameter manually measured for each mitochondrion.

To estimate the surface area of cristae, cristae were considered as flat disks with a diameter of $L_{2}$ (Supplementary figure 1A). Hence, the surface area of a crista is

$S_{crista}=2*\pi\left( \frac{L_{2}}{2} \right)^{2}$.

The surface area of all cristae was calculated in all analyzed mitochondria, where the length measured of each traced crista was used as $L_{2}$.

The [cristae/IMM]% of each mitochondrion was calculated by

$\left[ {cristae}/{IMM} \right]\%=\frac{S_{cristae}}{S_{cristae}+S_{boundary}}$.
